# Building tools for machine learning and artificial intelligence in cancer research: best practices and a case study with the PathML toolkit for computational pathology

**DOI:** 10.1101/2021.10.21.465212

**Authors:** Jacob Rosenthal, Ryan Carelli, Mohamed Omar, David Brundage, Ella Halbert, Jackson Nyman, Surya Narayanan Hari, Eliezer Mendel Van Allen, Luigi Marchionni, Renato Umeton, Massimo Loda

## Abstract

Imaging datasets in cancer research are growing exponentially in both quantity and information density. These massive datasets may enable derivation of insights for cancer research and clinical care, but only if researchers are equipped with the tools to leverage advanced computational analysis approaches such as machine learning and artificial intelligence. In this work, we highlight three themes to guide development of such computational tools: scalability, standardization, and ease of use. We then apply these principles to develop PathML, a general-purpose research toolkit for computational pathology. We describe the design of the PathML framework and demonstrate applications in diverse use-cases. PathML is publicly available at www.pathml.com.

## Big Data, image analysis, and machine learning in cancer research

Imaging has long been a cornerstone of cancer research and clinical care, providing insight into tissue morphology and spatial intercellular dynamics. Technological advances in recent years have enabled microscopy at a larger scale than ever before, leading to exponential growth in the size of commonly available datasets – a trend that is likely to continue to accelerate in coming years.

“Big Data” in biomedical imaging can be conceptualized along two orthogonal axes: sample size and data dimensionality (Figure 1). The first axis (*n*) can be measured by simply counting the number of cases in a dataset. Scaling in this dimension has been chiefly driven by advances in high-throughput imaging technologies. A notable example can be seen in the field of pathology, where increasing adoption of digital workflows results in slide scanning being routinely incorporated into pathologists’ workflows, consequently creating large databases of whole slide images (WSIs). Early adopters of digital pathology workflows are scanning more than 1 million slides per year [1] – several orders of magnitude larger than current benchmark datasets such as TCGA, and an indication of the potential volume of data that large academic tertiary care hospitals can expect to routinely generate as workflows are increasingly digitized.

**Figure 1.**
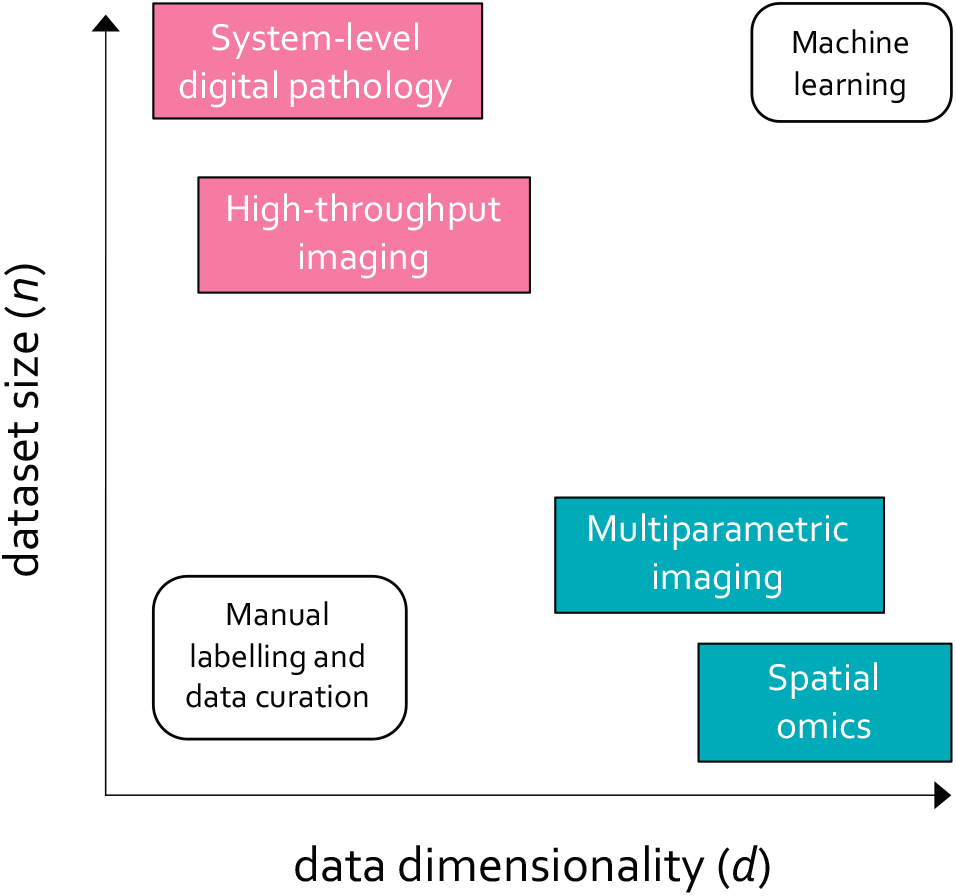
“Big Data” in biomedical imaging scales along two orthogonal axes: dataset size (*n*), which captures the number of data points (i.e. cases) in each dataset, and data dimensionality (*d*), which refers to the amount of data captured in each data point.

At the same time, data are also growing in the amount of information captured in each image, which we refer to as data dimensionality (*d*). This is chiefly driven by emerging technologies in spatial omics (i.e., spatial quantification of molecular markers such as proteins or RNA) and highly multiplexed imaging (reviewed in [2]). In contrast to brightfield images with three channels (red, green, and blue), each of these high-dimensional images may have upwards of 10,000 channels, each representing a specific target. Volumetric imaging further increases information content in each specimen by adding a depth dimension, enabling the capture of 3-dimensional tissue morphology. Thus, dataset sizes can grow even while the number of cases remains constant.

This rapid proliferation of imaging data has significant implications for cancer research, especially in conjunction with accompanying metadata such as genomics and outcomes. Large sample sizes provide sufficient power for discovery and quantification of histological patterns associated with clinically and biologically relevant features, with recent work demonstrating the potential of these methods to improve clinical and diagnostic workflows [3–5] and discover image-based biomarkers that recapitulate molecular features [6, 7]. Similarly, the rich contextual information captured in high-dimensional imaging data lays the groundwork for interrogation of tumor microenvironment at unprecedented resolution [8, 9]. The ubiquity of brightfield microscopy makes it an especially attractive candidate for image-based biomarker development, as digital workflows are increasingly deployed in a wider variety of clinical contexts.

However, while increasing scale of imaging datasets presents new opportunities and avenues of investigation, it also presents major challenges. Namely, these advances are only possible by leveraging computational image analysis methods, particularly deep learning. Deep learning models are flexible and powerful and have demonstrated remarkable success at identifying patterns in large datasets. As cancer research enters the age of “Big Data,” machine learning and is therefore poised to become an increasingly essential tool in the researcher’s toolkit, necessary for making use of massive datasets to study impactful questions in cancer biology and clinical care.

To enable this transition, software tools must lower the barrier to entry for computational image analysis, providing a bridge between the worlds of cancer research and machine learning. In this work, we discuss general features necessary for successful software tools, and present PathML: an open-source toolkit for computational pathology which we have built to address this outstanding need.

## Guiding principles for building software tools to accelerate research

To effectively leverage the wealth of imaging data, researchers must be equipped with the tools to easily incorporate powerful computational image analysis methods into their research. We identified three key elements that should guide design and development of software tools in this domain: scalability, standardization, and ease of use.

### Scalability

As datasets grow, analysis tools must be carefully designed to meet the technical challenges presented by scaling up in both *n* and *d*. Algorithms should be parallelized wherever possible, reducing computation time by running tasks concurrently. To enable efficient computation at the massive scale of tomorrow’s datasets, tools should embrace distributed processing and provide easy integration with commonly used open source big data solutions for on-premise, cloud, and hybrid infrastructures (e.g., Kubernetes, Hadoop YARN, Slurm, etc.). Support for hardware accelerators such as Graphics Processing Units (GPUs) and Tensor Processing Units (TPUs) is a requirement for computationally intensive tasks such as training machine learning models. Finally, tools should enable users to work with data that are larger than available memory – an important feature for accommodating larger data and supporting exploratory research on consumer-grade computers.

### Standardization

Another crucial consideration is standardization. No single tool can or should do everything; rather, by embracing standardized file formats, data structures, and Application Programming Interfaces (APIs), individual tools can focus on specialized tasks while still providing cross-compatibility with other tools. For example, researchers may need to implement domain-specific algorithms for working with specific data types of interest (e.g., stain deconvolution for H&E images), but should interface with industry-standard machine learning frameworks (e.g., PyTorch [10] and TensorFlow [11]) rather than implementing basic machine learning functionalities from scratch. In addition to providing consistency for users, this approach also promotes emergence of a cohesive ecosystem of tools, such as those built around the AnnData [12] standard in single-cell omics.

### Ease of use

A tool may be scalable and standardized, but it can only have an impact on accelerating research if users adopt it into their workflows. Therefore, software should be designed from the ground up with the intended audience in mind, and tools should be accessible with only minimal prior training in programming. This can be facilitated by building applications around well-defined APIs, which reduce the learning curve by providing consistency and by abstracting away some technical details from end users. All source code should be fully documented, with reproducible worked examples and detailed reference materials for all APIs. On the flip side of the coin, researchers will stand to benefit the most from advances in computational approaches if they are comfortable with the basics of coding in commonly used languages such as Python or R.

## PathML: a toolkit for computational pathology

We applied these guiding principles to development of PathML, an open-source toolkit designed for digital pathology research.

There are several existing tools that serve various needs in computational pathology. Some tools provide implementations of specific workflows or workflow components but are not designed as fully customizable, general-purpose libraries [13–17]. HistomicsTK [18] offers a Python API for running aspects of analysis workflows, but is built around a specific data management platform (Girder) rather than being platform-agnostic. QuPath [19] is an open-source tool for viewing and analyzing WSIs which has a scripting language for programmatic analysis but does not natively support Python. SquidPy [20] is primarily focused on spatial omics rather than general-purpose image analysis. There are also commercial tools available for digital pathology, some with support for machine learning analysis; however, these proprietary tools are not always ideal for researchers due to their cost and reduced flexibility in development relative to open source tools which enable full transparency into the underlying source code. Notably, there are no currently available open source tools which support the following requirements: the ability to load images from a wide array of file formats, including proprietary formats and standard formats such as TIFF and DICOM, under a common API; a standardized API for building custom preprocessing pipelines from modular components; support for running preprocessing at scale on commonly used high-performance computing solutions; integration with industry-standard machine learning frameworks in Python; and uniting analysis of brightfield and fluorescence images under a common framework.

To fill this unmet need, we developed PathML as a general-purpose toolkit for computational pathology, designed to be both highly performant and easy to use for researchers without requiring extensive training in programming or data science. PathML provides a general framework for creating and running preprocessing pipelines, unifying analysis of varying file formats (e.g., TIFF, DICOM, proprietary file formats from vendors, etc.), imaging modalities (e.g., H&E, IHC, Vectra Opal, CODEX, etc.), and dataset scales (from individual images to millions of images) under a single object-oriented API, with data structures and design choices specifically tailored to digital pathology (Figure 2). The PathML library is written in Python 3 to promote ease of use and integration with the broader ecosystem of standard tools for data science and machine learning; however, we leverage libraries such as NumPy[21] and PyTorch which are written in low-level languages such as C, C++, and CUDA to handle computationally intensive operations more efficiently. An extensive suite of unit testing and integration testing helps ensure that all code in PathML is free of bugs and is working as expected. By developing PathML as an open source tool, we hope to build a community of users and collaborators to collectively accelerate the pace of innovation in digital pathology research.

**Figure 2.**
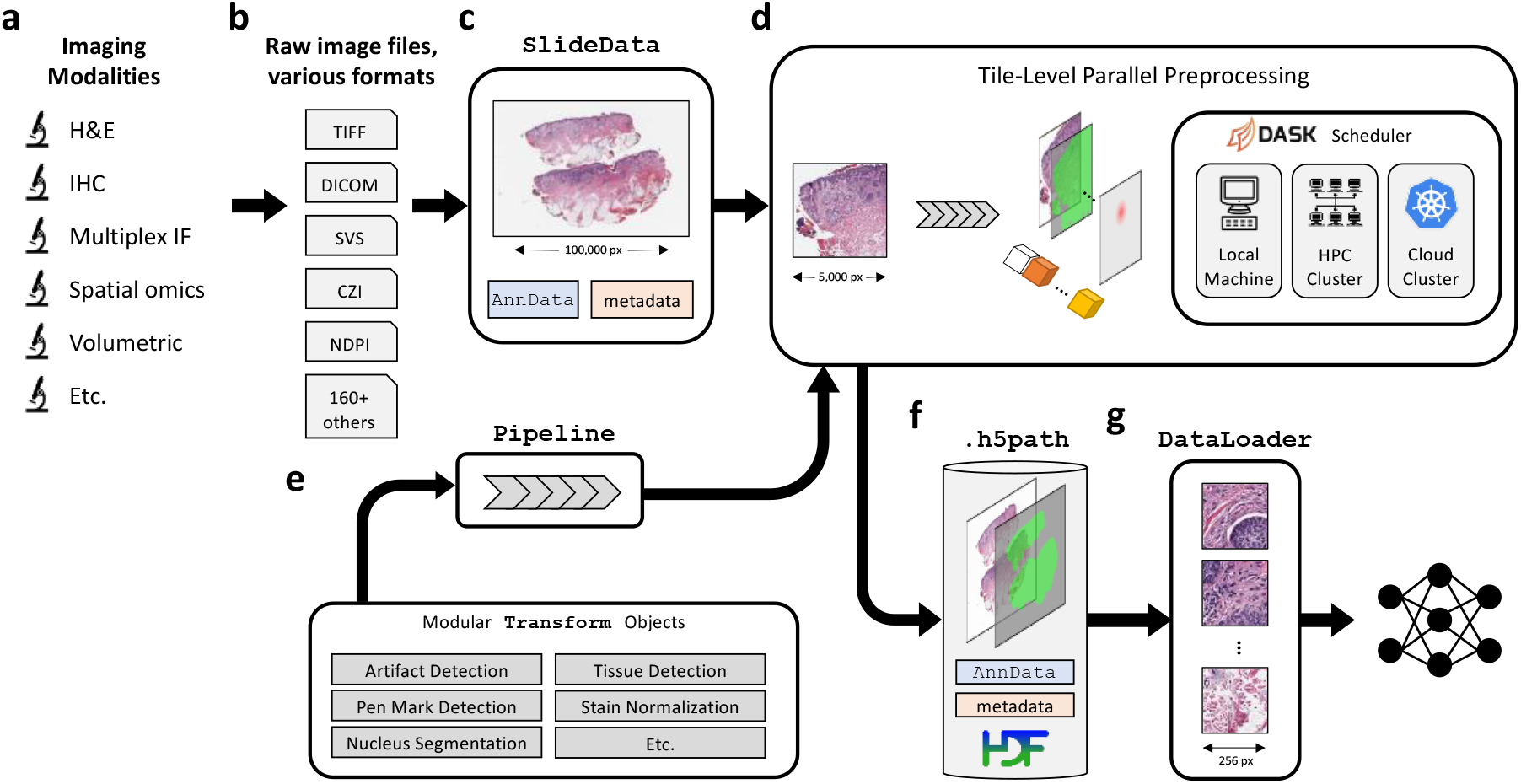
Overview of PathML preprocessing framework. (**a**) A wide range of imaging platforms and modalities are supported via (**b**), support for loading a comprehensive set of more than 165 file formats, including proprietary formats from vendors (see full list of supported file formats in Supplementary Table 1). (**c**) Raw image files are loaded into SlideData objects, which encapsulate the image as well as associated metadata. (**d**) To enable efficient processing of gigapixel-scale scans, images are divided into tiles and preprocessing pipelines are applied independently to each tile. Tiles can thus be processed in parallel, using the Dask scheduler to orchestrate distributed computation on large clusters, with support for both cloud and on-premise computing. Smaller images are processed in this framework as a single tile containing the entire image. (**e**) A preprocessing pipeline is defined as a set of transformations applied sequentially. Transformations are modular, so can be mix-and-matched to rapidly build custom pipelines. (**f**) Processed tiles are aggregated into an h5path file on disk, along with associated metadata such as labels, masks, and counts matrix. Hierarchical Data Format (HDF5) is used to enable efficient slicing and indexing of the resulting file without needing to load the entire file into memory. (**g**) DataLoaders from frameworks such as PyTorch then interact with the h5path file to efficiently feed images from the processed image into downstream machine learning models.

The first step in a PathML workflow is loading the raw image file to create a SlideData object, which is the central data class representing an image and associated metadata. To accommodate the wide array of file formats commonly used in digital pathology, we provide three separate backends for reading image files, each supporting a complementary set of file formats (Supplementary Table 1). Each backend adheres to a standardized API, enabling users to manipulate images with a consistent interface regardless of file format or imaging modality (Supplementary Vignette 1).

The next step is to create a preprocessing pipeline, which we define as the sequential application of independent building blocks, or transformations. Each transformation applies a specific operation which may include modifying an input image, creating or modifying pixel-level metadata (i.e., masks), or creating or modifying image-level metadata (e.g., image quality metrics or an AnnData counts matrix). Transformations are general and flexible, providing a standardized interface to compose preprocessing pipelines. We provide in PathML a set of commonly used transformations, both domain-specific (e.g., H&E stain deconvolution, tissue detection, WSI artifact detection) and general-purpose (e.g., blurring, binary thresholding) (Supplementary Figure 1). Users may also implement custom transformations, and we provide an API to enable integration of custom transformations alongside pre-built transformations. Multiple transformations can be composed into a single compound transformation. Transformations therefore provide the building blocks for formalizing the design and implementation of arbitrary preprocessing pipelines. This API allows researchers to write scalable, end-to-end preprocessing pipelines in only a few lines of code, using the same syntax and building blocks across different file formats and imaging modalities (Supplementary Vignettes 2 and 3).

One of the most common technical challenges in computational pathology is presented by extremely large file sizes, with high-resolution WSIs routinely exceeding the capacity of available memory. We therefore designed PathML based on a paradigm of independent processing of tiles. To run a preprocessing pipeline, subregions of the image (i.e., tiles) are extracted and passed to the preprocessing pipeline independently. Smaller images are processed in this framework as a single tile containing the entire image. All processed tiles are then aggregated together into an on-disk array optimized for storing and manipulating large imaging datasets. This design allows for efficient preprocessing of large datasets of gigapixel images, as the data parallelism approach can efficiently scale up to make use of additional computational resources (e.g., cores in a multi-core computing unit, computing nodes in a cluster, etc.). We use the dask.distributed[22] scheduler on the backend, which allows for distributed preprocessing on many common high-performance computing platforms, including support for both on-premise and cloud computing environments. Importantly, tile extraction and distributed processing are handled automatically by PathML, enabling users to leverage these features to run analyses at scale with no change to the rest of their code. One limitation of this tile-centric approach is that artifacts may arise when tiles are aggregated back together, such as discontinuities at tile edges or “patchwork” effects. However, processing is inherently limited by the number of pixels that can be stored and manipulated in memory at once, so there is always a tradeoff between processing few low-resolution tiles and many high-resolution tiles. Users have complete control over tile extraction parameters, including the ability to use overlapping tiles which, in conjunction with stitching algorithms such as [23] can mitigate such artifacts.

As tiles are processed, they are aggregated together and written to disk. We define a file specification (h5path) which leverages the Hierarchical Data Format (HDF5) to enable efficient read/write access to regions of the processed image without loading the entire image into memory. Along with the processed images and masks, each h5path file contains associated slide-level and tile-level metadata. Each SlideData object is backed by a corresponding h5path file on disk, allowing for intuitive object-oriented workflows scalable to larger-than-memory images.

After a preprocessing pipeline has been run, we provide utilities to load the processed images into machine learning frameworks for downstream tasks (e.g., PyTorch DataLoaders). Preprocessing pipelines may themselves include transformations which encapsulate machine learning algorithms, for example using a model to perform nucleus detection and/or classification on each tile. PathML further provides PyTorch implementations of commonly used models such as U-Net [24] and HoVer-Net [25]. Finally, we provide streamlined access to domain-specific datasets including PanNuke [26], PESO [27], and DeepFocus [28] for use in model training and benchmarking for various tasks. With support from open-source contributors, we hope that the inventory of available datasets and machine learning models will continue to expand.

In sum, PathML provides comprehensive support for each step in the computational pathology research workflow. We define a framework for preprocessing images and metadata which is streamlined and flexible for a wide variety of file formats and imaging modalities, implemented in an efficient, open source, fully tested and thoroughly documented Python package. We have already applied PathML to enable published [29] and currently ongoing computational pathology research at our institutions; by releasing it as an open source standard toolkit to bridge the gap between digital pathology and the broader machine learning and artificial intelligence ecosystem, we aim to lower the barrier to entry and accelerate progress in digital pathology research, thus benefiting the entire research community and moving one step closer to implementation of computational methods in the clinic.

## Conclusion

With biomedical imaging datasets growing exponentially in both number of samples and dimensionality (i.e., data within each sample), machine learning is emerging as an increasingly essential tool for cancer researchers. To support these efforts, software tools must be designed with emphasis on scalability, standardization, and ease of use. Here we introduce PathML, a framework built with these best practices in mind that aims at lowering the barrier of entry to digital pathology, and show how a number of heterogeneous computational pathology use cases can be readily implemented in very few lines of code. With comprehensive support for all aspects of computational pathology research, from loading a wide variety of imaging modalities and file formats, to building modular and completely customizable pre-processing pipelines, to parallel-computing provisions, and integrations with other tools in the machine learning, AI, and single-cell analysis ecosystems, PathML can be employed to tackle a variety of biologically relevant problems. We anticipate that the real impactful part of this work will be around the applications of this technology, which we made open-source and therefore available to all researchers. PathML is publicly available at www.pathml.com.

## Supporting information

Supplementary Vignette 1

Supplementary Vignette 2

Supplementary Vignette 3

## Acknowledgements

The authors would like to thank Jason Johnson, Jerri Zhang, the Artificial Intelligence Operations and Data Science Services group, and many collaborators in the Department of Informatics & Analytics at Dana-Farber Cancer Institute and in the Department of Pathology and Laboratory Medicine at Weill Cornell Medicine for their continuous support and critical feedback that improved this work. We thank Angeles Duran, Jorge Moscat, and Maria T. Diaz-Meco for providing the images used in Supplementary Figure 1, panels f and g. ML’s work is supported by NCI P50CA211024, DoD PC160357, DoD PC180582, and the Prostate Cancer Foundation.

## Supplementary Material

**Supplementary Figure 1.**
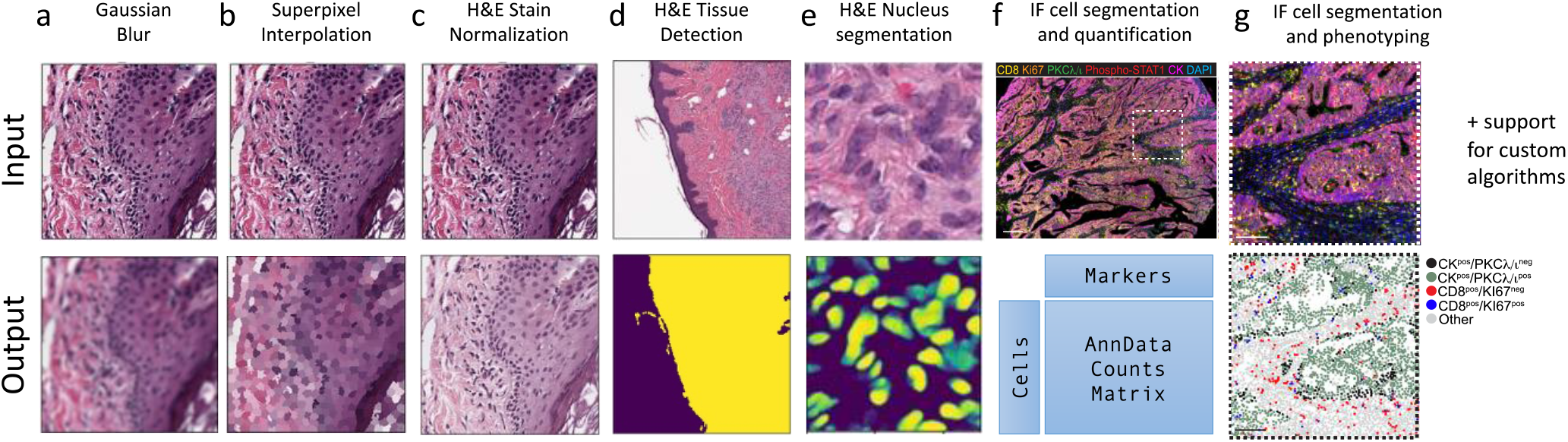
A preprocessing pipeline is defined as a sequence of modular transformations. Here, we show selected examples to demonstrate the generalizability and flexibility of the PathML preprocessing API: (**a**) Gaussian blur, (**b**) superpixel interpolation using the algorithm described in [30], (**c**) H&E stain normalization using the method described in [31], (**d**) H&E tissue detection using the method described in [32], (**e**) H&E nucleus segmentation using a pre-trained HoVer-Net neural network model [25], (**f**) immunofluorescence cell segmentation and quantification using the AnnData standard data format [12], and (**g**) immunofluorescence cell segmentation and rules-based phenotyping of subregion outlined in (f) (from [29]). Each transformation is parametrized by one or more variables, allowing users full control to modulate the outputs at each step (not shown). Additionally, the modular API allows researchers to mix-and-match pre-made transformations alongside custom operations, enabling streamlined construction of completely customizable preprocessing pipelines.

**Supplementary Table 1.**
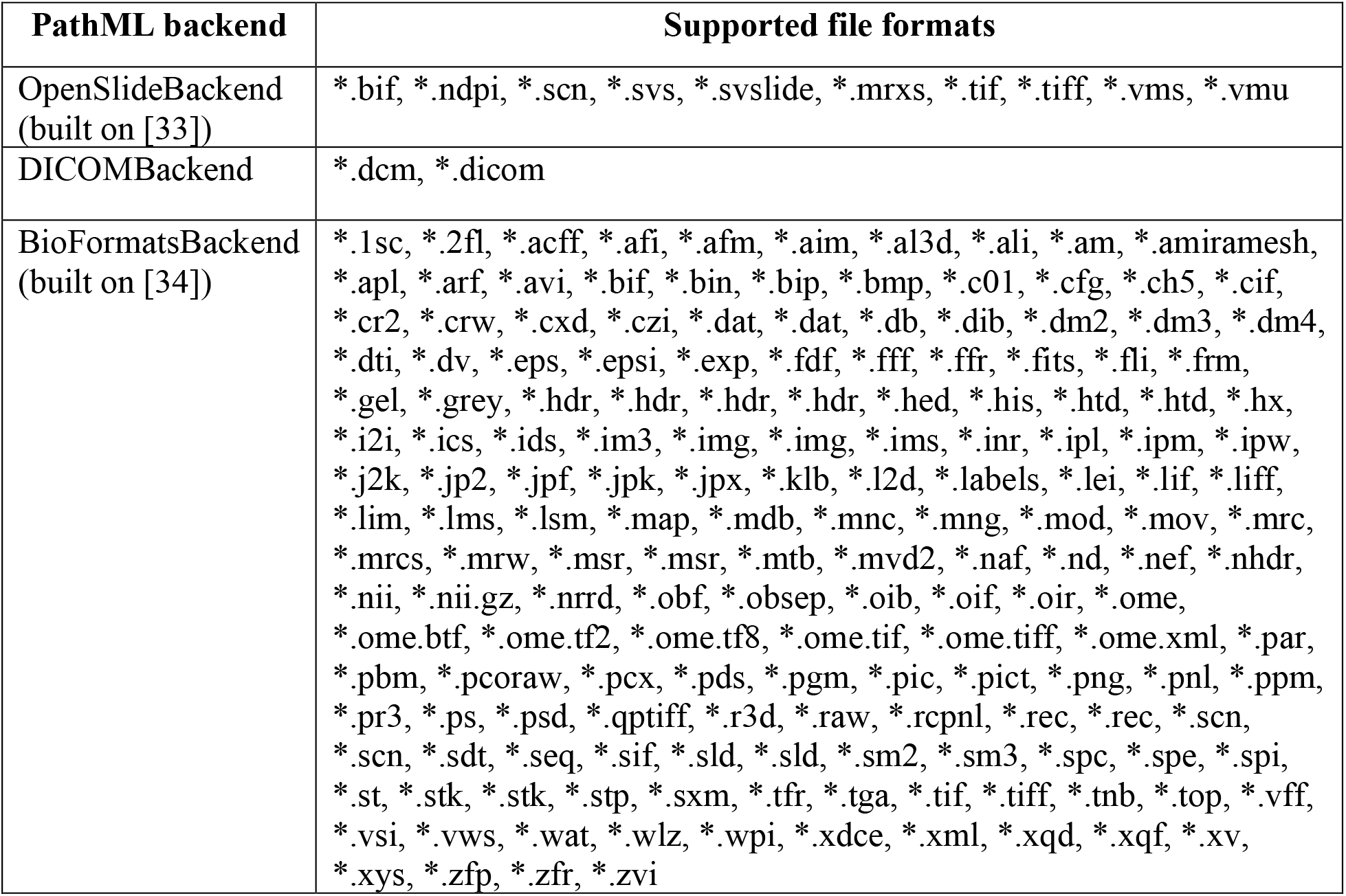
List of file formats currently supported by PathML through its three backends. Each backend is modular and built on a consistent API, providing a consistent interface for users regardless of file format and facilitating implementation of additional backends to support other file types as needed. See also Supplementary Vignette 1 for more detailed code snippets highlighting the PathML API for loading images.

