## Supplementary Vignette 1 for "Building tools for machine learning and artificial intelligence in cancer research: best practices and a case study with the PathML toolkit for computational pathology"

loading\_images\_vignette


### Supplementary Vignette 1¶

#### Consistent API for loading images from diverse modalities and file formats¶

`PathML` provides support for loading a wide array of imaging modalities and file formats under a standardized syntax.

In this vignette, we highlight code snippets for loading a range of image types ranging from brightfield H&E and IHC to highly multiplexed immunofluorescence and spatial expression and proteomics, from small images to gigapixel scale:

| Imaging modality | File format | Source | Image dimensions (X, Y, Z, C, T) |
| --- | --- | --- | --- |
| Brightfield H&E | Aperio SVS | OpenSlide example data | (32914, 46000, 1, 3, 1) |
| Brightfield H&E | Generic tiled TIFF | OpenSlide example data | (32914, 46000, 1, 3, 1) |
| Brightfield IHC | Hamamatsu NDPI | OpenSlide example data | (73728, 126976, 1, 3, 1) |
| Brightfield H&E | Hamamatsu VMS | OpenSlide example data | (76288, 102400, 1, 3, 1) |
| Brightfield H&E | Leica SCN | OpenSlide example data | (153470, 53130, 1, 3, 1) |
| Fluorescence | MIRAX | OpenSlide example data | (170960, 76324, 1, 3, 1) |
| Brightfield IHC | Olympus VSI | OpenSlide example data | (6753, 13196, 1, 3, 1) |
| Brightfield H&E | Trestle TIFF | OpenSlide example data | (25408, 61504, 1, 3, 1) |
| Brightfield H&E | Ventana BIF | OpenSlide example data | (93951, 105813, 1, 3, 1) |
| Fluorescence | Zeiss ZVI | OpenSlide example data | (1388, 1040, 13, 3, 1) |
| Brightfield H&E | DICOM | Orthanc example data | (30462, 78000, 1, 3, 1) |
| Fluorescence (CODEX spatial proteomics) | TIFF | Schurch et al., Cell 2020 | (1920, 1440, 17, 4, 23) |
| Fluorescence (time-series + volumetric) | OME-TIFF | OME-TIFF example data | (512, 512, 10, 2, 43) |
| Fluorescence (MERFISH spatial gene expression) | TIF | Zhuang et al., 2020 | (2048, 2048, 7, 1, 40) |
| Fluorescence (Visium 10x spatial gene expression) | TIFF | 10x Genomics | (25088, 26624, 1, 1, 4) |

All images used in these examples are publicly available for download at the links listed above.

Note that across the wide diversity of modalities and file formats, the syntax for loading images is consistent (see examples below).

In [1]:

```
# import utilities for loading images
from pathml.core import HESlide, CODEXSlide, VectraSlide, SlideData, types
```

##### Aperio SVS¶

In [2]:

```
my_aperio_image = HESlide("./data/CMU-1.svs")
```

##### Generic tiled TIFF¶

In [3]:

```
my_generic_tiff_image = HESlide("./data/CMU-1.tiff", backend = "bioformats")
```

##### Hamamatsu NDPI¶

The `labels` field can be used to store slide-level metadata. For example, in this case we store the target gene, which is Ki-67:

In [4]:

```
my_ndpi_image = SlideData("./data/OS-2.ndpi", 
                          labels = {"taget" : "Ki-67"},
                          slide_type = types.IHC)
```

##### Hamamatsu VMS¶

In [5]:

```
my_vms_image = HESlide("./data/CMU-1/CMU-1-40x - 2010-01-12 13.24.05.vms", backend = "openslide")
```

##### Leica SCN¶

In [6]:

```
my_leica_image = HESlide("./data/Leica-1.scn")
```

##### MIRAX¶

In [7]:

```
my_mirax_image = SlideData("./data/Mirax2-Fluorescence-1/Mirax2-Fluorescence-1.mrxs", 
                           slide_type = types.IF)
```

##### Olympus VSI¶

Again, we use the `labels` field to store slide-level metadata such as the name of the target gene.

In [8]:

```
my_olympus_vsi = SlideData("./data/OS-3/OS-3.vsi", 
                           labels = {"taget" : "PTEN"},
                           slide_type = types.IHC)
```

##### Trestle TIFF¶

In [9]:

```
my_trestle_tiff = SlideData("./data/CMU-2/CMU-2.tif")
```

##### Ventana BIF¶

In [10]:

```
my_ventana_bif = SlideData("./data/OS-1.bif")
```

##### Zeiss ZVI¶

Again, we use the `labels` field to store slide-level metadata such as the name of the target gene.

In [11]:

```
my_zeiss_zvi = SlideData("./data/Zeiss-1-Stacked.zvi", 
                         labels = {"target" : "HER-2"},
                         slide_type = types.IF)
```

##### DICOM¶

In [12]:

```
my_dicom = HESlide("./data/orthanc_example.dcm")
```

##### Volumetric + time-series OME-TIFF¶

In [13]:

```
my_volumetric_timeseries_image = SlideData(
    "./data/tubhiswt-4D/tubhiswt_C1_TP42.ome.tif", 
    labels = {"organism" : "C elegans"},
    volumetric = True,
    time_series = True,
    backend = "bioformats"
)
```

##### CODEX spatial proteomics¶

The `labels` field can be used to store whatever slide-level metadata the user wants; here we specify the tissue type

In [14]:

```
my_codex_image = CODEXSlide('../../data/reg031_X01_Y01.tif', 
                            labels = {"tissue type" : "CRC"});
```

##### MERFISH spatial gene expression¶

In [15]:

```
my_merfish_image = SlideData("./data/aligned_images0.tif", backend = "bioformats")
```

##### Visium 10x spatial gene expression¶

Here we load an image with accompanying expression data in `AnnData` format.

In [16]:

```
# load the counts matrix of spatial genomics information
import scanpy as sc
adata = sc.read_10x_h5("./data/Visium_FFPE_Mouse_Brain_IF_raw_feature_bc_matrix.h5") 

# load the image, with accompanying counts matrix metadata
my_visium_image = SlideData("./data/Visium_FFPE_Mouse_Brain_IF_image.tif", 
                            counts=adata, 
                            backend = "bioformats")
```

```
Variable names are not unique. To make them unique, call `.var_names_make_unique`.
Variable names are not unique. To make them unique, call `.var_names_make_unique`.
```

##### Summary¶

The `PathML` API provides a consistent, easy to use interface for loading a wide range of imaging data:

- from standard file formats to vendor-specific proprietary file formats
- from small images to gigapixel-scale images
- from brightfield to fluorescence
- etc.

The output from all of the code snippets above is a `SlideData` object compatible with the `PathML` preprocessing module.

Full documentation of the `PathML` API is available at https://pathml.org.

Full code for this vignette is available at https://github.com/Dana-Farber-AIOS/pathml/tree/master/examples/manuscript\_vignettes\_stable
