## Supplementary Vignette 2 for "Building tools for machine learning and artificial intelligence in cancer research: best practices and a case study with the PathML toolkit for computational pathology"

workflow\_HE\_vignette


### Supplementary Vignette 2¶

#### Example workflow for H&E images¶

Here we demonstrate a typical workflow for preprocessing of H&E images, consisting of the following steps:

1. Loading the raw image
2. Defining a simple preprocessing pipeline for tissue detection
3. Creating a PyTorch DataLoader for interfacing with any downstream machine learning model

The image used in this example is publicly avilalable for download: http://openslide.cs.cmu.edu/download/openslide-testdata/Aperio/

**a. Load the image**

In [3]:

```
from pathml.core import SlideData, types

# load the image
wsi = SlideData("../../data/CMU-1.svs", name = "example", slide_type = types.HE)
```

**b. Define a preprocessing pipeline**

Pipelines are created by composing a sequence of modular transformations; in this example we apply a blur to reduce noise in the image followed by tissue detection

In [5]:

```
from pathml.preprocessing import Pipeline, BoxBlur, TissueDetectionHE

pipeline = Pipeline([
    BoxBlur(kernel_size=15),
    TissueDetectionHE(mask_name = "tissue", min_region_size=500, 
                      threshold=30, outer_contours_only=True)
])
```

**c. Run preprocessing**

Now that we have constructed our pipeline, we are ready to run it on our WSI.
PathML supports distributed computing, speeding up processing by running tiles in parallel among many workers rather than processing each tile sequentially on a single worker. This is supported by Dask.distributed on the backend, and is highly scalable for very large datasets.

The first step is to create a `Client` object. In this case, we will use a simple cluster running locally; however, Dask supports other setups including Kubernetes, SLURM, etc. See the PathML documentation for more information.

In [6]:

```
from dask.distributed import Client, LocalCluster

cluster = LocalCluster(n_workers=6)
client = Client(cluster)

wsi.run(pipeline, distributed=True, client=client);
```

In [7]:

```
print(f"Total number of tiles extracted: {len(wsi.tiles)}")
```

```
Total number of tiles extracted: 150
```

**e. Save results to disk**

The resulting preprocessed data is written to disk, leveraging the HDF5 data specification optimized for efficiently manipulating larger-than-memory data.

In [8]:

```
wsi.write("./data/CMU-1-preprocessed.h5path")
```

**f. Create PyTorch DataLoader**

The `DataLoader` provides an interface with any machine learning model built on the PyTorch ecosystem

In [9]:

```
from pathml.ml import TileDataset
from torch.utils.data import DataLoader

dataset = TileDataset("./data/CMU-1-preprocessed.h5path")
dataloader = DataLoader(dataset, batch_size = 16, num_workers = 4)
```

##### Summary¶

Here we demonstrate a complete `PathML` workflow for analyzing brightfield images:

1. Loading the raw image
2. Define a simple preprocessing pipeline for tissue detection
3. Create a PyTorch DataLoader for interfacing with any downstream machine learning model

Full documentation of the `PathML` API is available at https://pathml.org.

Full code for this vignette is available at https://github.com/Dana-Farber-AIOS/pathml/tree/master/examples/manuscript\_vignettes\_stable
