## Supplementary Vignette 3 for "Building tools for machine learning and artificial intelligence in cancer research: best practices and a case study with the PathML toolkit for computational pathology"

workflow\_IF\_vignette


### Supplementary Vignette 3¶

#### Example workflow for immunofluorescence images¶

Here we demonstrate a typical workflow for preprocessing of immunofluorescence images, comsisting of the following steps:

1. Loading a raw image in TIFF format
2. Defining a preprocessing pipeline for cell segmentation and marker quantification for each cell
3. Integrating with other commonly used tools such as `Scanpy` for working with the quantified cell-level data:
   - dimensionality reduction
   - clustering
   - co-occurence analysis
   - visualization

The image used in this example is a tissue microarray (TMA) generated on the CODEX spatial proteomics imaging platform, from Schurch et al., *Coordinated Cellular Neighborhoods Orchestrate Antitumoral Immunity at the Colorectal Cancer Invasive Front* (Cell, 2020). The image used in this example is publicly avilalable for download from the Cancer Imaging Archive: https://doi.org/10.7937/tcia.2020.fqn0-0326

**a. Load the image**

In [2]:

```
from pathml.core.slide_data import CODEXSlide

# load the image
slidedata = CODEXSlide('../../data/reg031_X01_Y01.tif', 
                       labels = {"tissue type" : "CRC"});
```

The CODEX imaging protocol is cyclic, so markers are imaged in groups of 4.
These images use the standard convention of (X, Y, Z, C, T) channel order.
In this case, the time dimension (T) is being used to denote cycles; here we see that the image has 17 z-slices for 23 cycles of 4 markers each.

In [3]:

```
# These tif are of the form (x,y,z,c,t) but t is being used to denote cycles
# 17 z-slices, 4 channels per 23 cycles, 70 regions
slidedata.slide.shape
```

Out[3]:

```
(1920, 1440, 17, 4, 23)
```

**b. Define a preprocessing pipeline**

Pipelines are created by composing a sequence of modular transformations; in this example we first choose a z-slice from our CODEX image, then segment the cells using the pre-trained Mesmer machine learning model, and finally quantify the expression of each protein in each cell.

In [4]:

```
from pathml.preprocessing import (
    Pipeline, CollapseRunsCODEX, SegmentMIF, QuantifyMIF
)

# 31 -> Na-K-ATPase
pipeline = Pipeline([
    CollapseRunsCODEX(z=6),
    SegmentMIF(model='mesmer', nuclear_channel=0, cytoplasm_channel=31, image_resolution=0.377442),
    QuantifyMIF(segmentation_mask='cell_segmentation')
])
```

**c. Run preprocessing**

In this example, we choose not to distribute computation as the image is relatively small. Instead, we process the image as a single tile:

In [ ]:

```
slidedata.run(pipeline, distributed=False, tile_size=slidedata.shape);
```

In [6]:

```
print(f"Total number of tiles extracted: {len(slidedata.tiles)}")
```

```
Total number of tiles extracted: 1
```

**e. Save results to disk**

The resulting preprocessed data is written to disk, leveraging the HDF5 data specification optimized for efficiently manipulating larger-than-memory data.

In [7]:

```
slidedata.write("./data/reg031_X01_Y01-preprocessed.h5path")
```

**f. AnnData Integration and Spatial Single Cell Analysis**

Now let's explore the single-cell quantification of our imaging data. Our pipeline produced a single-cell matrix of shape (n\_cell x n\_proteins) where each cell has attached additional information including location on the slide and the size of the cell in the image. This information is stored in `slidedata.counts` as an `Anndata` object (https://anndata.readthedocs.io/en/latest/anndata.AnnData.html).

In [8]:

```
adata = slidedata.counts.to_memory()
```

In [9]:

```
adata
```

Out[9]:

```
AnnData object with n_obs × n_vars = 2925 × 92
    obs: 'x', 'y', 'coords', 'filled_area', 'slice', 'euler_number', 'tile'
    obsm: 'spatial'
    layers: 'min_intensity', 'max_intensity'
```

`AnnData` is a standard data format, so this `AnnData` object gives us access to the entire Python (or Seurat) ecosystem of single cell analysis tools. We follow a single cell analysis workflow described in https://scanpy-tutorials.readthedocs.io/en/latest/pbmc3k.html and https://www.embopress.org/doi/full/10.15252/msb.20188746.

First we look at a violin plot for three randomly selected markers:

In [10]:

```
import scanpy as sc
sc.pl.violin(adata, keys = ['0','24','60'])
```

Next, we use UMAP to look at the cells in a low-dimensional visualization, colored by expression levels of the three markers:

In [11]:

```
sc.pp.log1p(adata)
sc.pp.scale(adata, max_value=10)
sc.tl.pca(adata, svd_solver='arpack')
sc.pp.neighbors(adata, n_neighbors=10, n_pcs=10)
sc.tl.umap(adata)
sc.pl.umap(adata, color=['0','24','60'])
```

```
OMP: Info #271: omp_set_nested routine deprecated, please use omp_set_max_active_levels instead.
```

Here, we perform Leiden clustering in the expression space:

In [12]:

```
sc.tl.leiden(adata, resolution = 0.15)
sc.pl.umap(adata, color='leiden')
```

Next, we use a dotplot to visualize the markers for each group:

In [13]:

```
sc.tl.rank_genes_groups(adata, 'leiden', method='t-test')
sc.pl.rank_genes_groups_dotplot(adata, groupby='leiden', vmax=5, n_genes=5)
```

```
WARNING: dendrogram data not found (using key=dendrogram_leiden). Running `sc.tl.dendrogram` with default parameters. For fine tuning it is recommended to run `sc.tl.dendrogram` independently.
```

Here, we plot the clustering results in the spatial domain, highlighting the spatial organization of the tissue:

In [14]:

```
sc.pl.spatial(adata, color='leiden', spot_size=15)
```

Finally, we compute the co-occurrence probability of the clusters, highlighting the interface with spatial analysis tools such as Squidpy: https://github.com/theislab/squidpy.

In [15]:

```
import squidpy as sq
sq.gr.co_occurrence(adata, cluster_key="leiden")
sq.pl.co_occurrence(
    adata,
    cluster_key="leiden"
)
```

##### Summary¶

Here we demonstrate a complete `PathML` workflow for analyzing immunofluorescence images:

1. Loading a raw image in TIFF format
2. Defining a preprocessing pipeline for cell segmentation and marker quantification for each cell
3. Integrating with other commonly used tools such as `Scanpy` for working with the quantified cell-level data:
   - dimensionality reduction
   - clustering
   - co-occurence analysis
   - visualization

Full documentation of the `PathML` API is available at https://pathml.org.

Full code for this vignette is available at https://github.com/Dana-Farber-AIOS/pathml/tree/master/examples/manuscript\_vignettes\_stable
